# reCRAC: A Stringent Method for Precise Mapping of Protein-RNA Interactions in Yeast

**DOI:** 10.1101/2024.05.22.594286

**Authors:** Michaela Ristova, Vadim Shchepachev, David Tollervey

## Abstract

Intricate interactions between RNA-binding proteins (RBPs) and RNA play pivotal roles in cellular homeostasis, impacting a spectrum of biological processes vital for survival. UV crosslinking methods to study protein-RNA interactions have been instrumental in elucidating their interactions but can be limited by degradation of target proteins during the process, low signal-to-noise ratios, and non-specific interactions. Addressing these limitations, we describe reCRAC (reverse CRAC), a novel adaptation of the CRAC (crosslinking and analysis of cDNA) technique, optimized for yeast *Saccharomyces cerevisiae*. Like CRAC, reCRAC applies tandem affinity purification to yield highly enriched protein preparations. However, reCRAC is redesigned to work with unstable proteins. This is achieved by lysing the cells directly into highly denaturing buffer conditions, followed by stringent purification steps. The reCRAC method was successfully applied to the easily degraded yeast protein Pin4, allowing identification of precise binding sites at base-pair resolution with greatly reduced target protein degradation and improved signal-to-noise ratios.

## 1. Introduction

RNA-binding proteins (RBPs) are critical to cellular function, playing crucial roles across a vast array of biological processes. These include transcription, as well as a host of post-transcriptional activities such as RNA processing, splicing, nuclear export, subcellular localization, and the precise control of mRNA stability, translation, and eventual decay ^1,2^. By adeptly facilitating necessary changes in the transcriptome in response to environmental stimuli, RBPs rapidly modulate gene expression, making them instrumental in maintaining cellular homeostasis, and ultimately ensuring the survival of the organism. In recent years, methodological advancements have transformed our ability to explore protein-RNA interactions within their native cellular environment, offering novel insights into the dynamic and often transient encounters between RNAs and protein partners. UV crosslinking lies at the heart of most of these methods ^3^. This zero-distance crosslinking initiates covalent bonds between directly interacting RNA and proteins, preserving the interactions between protein and RNAs even under very stringent extraction conditions ^4^. This represents a notable advance over techniques based on formaldehyde crosslinking, which trigger undesired protein-protein crosslinks, are less penetrative for large complexes, and the protein-RNA interactions can only be purified under semi-denaturing conditions, potentially generating false positive interactors ^5–7^.

Several protein-centric techniques to enrich the target proteins with the crosslinked RNAs have been reported, which differ in the method of library preparation ^5^. Commonly used techniques, eCLIP, iCLIP, irCLIP, easy CLIP, and CRAC utilize 254 nm UVC light to induce crosslinking ^8^. In contrast, PAR-CLIP employs 365 nm UVB light to exclusively crosslink RNA incorporating the uracil analogue 4-thioUracil (4sU), achieving a higher specificity and allowing for the study of dynamic processes such as RNA synthesis and turnover ^9^. Non-radioactive, fluorescence-based CLIP versions are also available ^10^. Subsequent protein purification is performed under varying conditions, from semi-denaturing (CLIP) to fully denaturing (CRAC).

RNA-centric techniques recover bound proteins and include silica-based methods such as 2C, TRAPP, and PAR-TRAPP, or phase-separation techniques like PTex, OOPS, and XRNAX ^11–13^. All of these techniques are reviewed by Esteban-Serna et al. ^14^.

This paper presents reCRAC (**re**verse **CRAC**), a novel modification of the tandem affinity based CRAC (crosslinking and analysis of cDNA) method applied in yeast ^15,16^. reCRAC offers a more robust approach for the isolation of protein-RNA complexes by decreasing target protein degradation, and therefore, enhancing the detection of authentic RNA targets. It achieves this by implementing more stringent and denaturing conditions, particularly during cell lysis and the first round of affinity purification, relative to CRAC and CLIP protocols. This protects the target protein from proteases degradation, decreases non-specific interactions, and minimizes background noise. The cell lysis buffer includes 6 M Guanidine-HCl and 300 mM salt, and is followed by denaturing purification on the Nickel (Ni-NTA) beads. The subsequent secondary purification is on anti-Flag beads, reversing the order of tandem purification steps in the CRAC method.

Briefly, reCRAC starts by engineering yeast strains to express the protein of interest tagged with a Flag-4xAla-8xHis N-terminal sequence (N-FH), or 8xHis-4xAla-Flag C-terminal sequence (HF-C) from the endogenous gene locus. This ensures that the fusion protein, regulated by its natural promoter, is the only variant produced. Strains in growth medium are irradiated with UVC light at 254 nm to facilitate formation of covalent bonds between the target proteins and interacting RNAs. After denaturing cell lysis, complexes are bound to Nickel Ni-NTA affinity resin. Enzymatic processing is then carried out on the Ni-NTA column including the partial RNase digestion, radioactive labelling with [^32^P], and linker ligation on the 3’ and 5’ ends of RNA. Following elution from the Nickel column, a second purification on anti-Flag beads is performed under more native conditions that are permissive for antibodies. Protein-RNA complexes are eluted, run on denaturing SDS-PAGE, and the RNA can is visualised by exposing the gel to film. The target protein-RNA complexes are excised from the gel and treated with Proteinase K to release an RNA fragment including the protein binding site. This RNA is recovered, converted to cDNA through reverse transcription, PCR amplified, and sequenced. The outline of the experimental steps in reCRAC and their approximate time duration is given in **Figure 1**.

**Figure 1.**
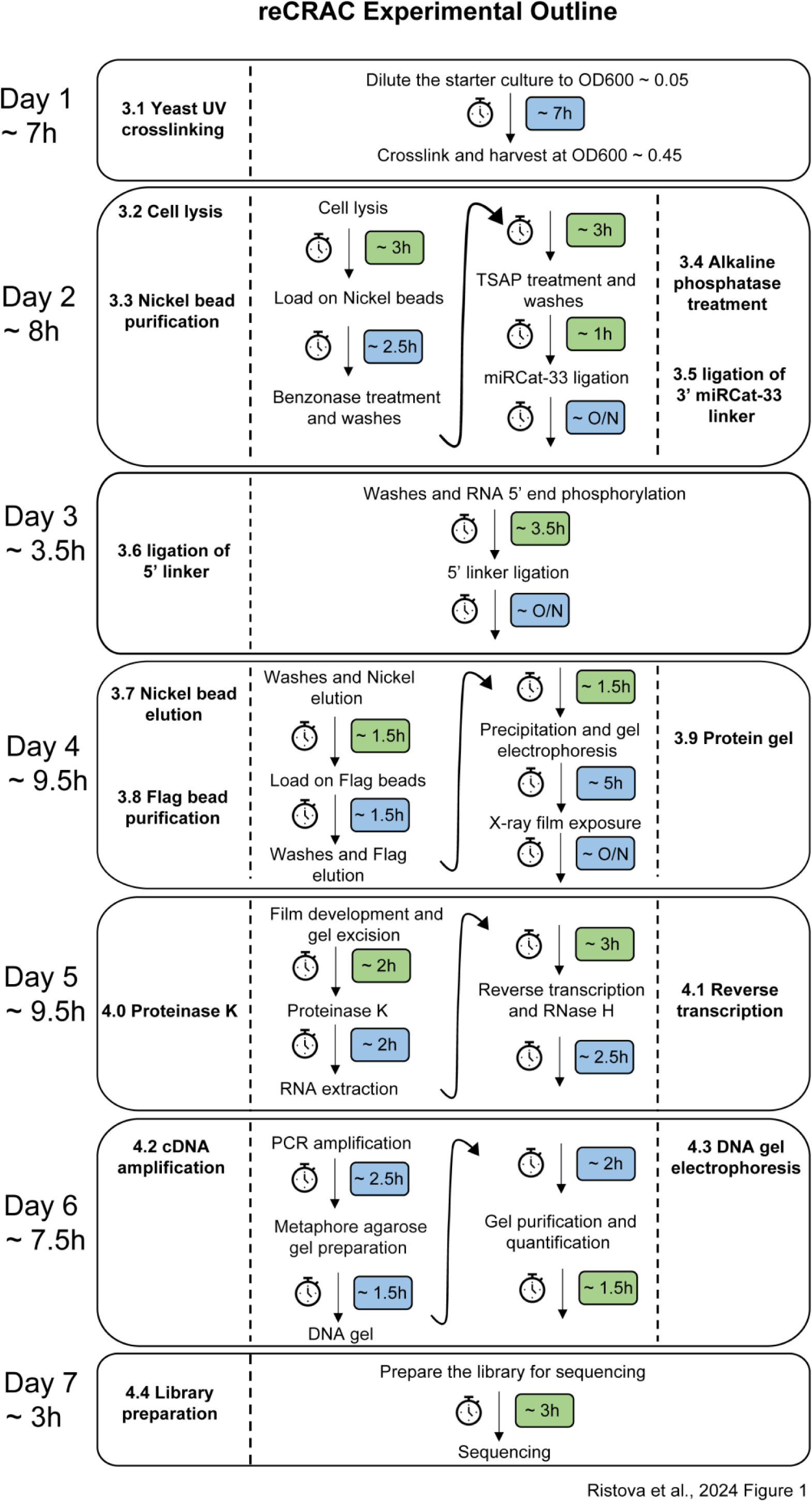
Outline of the reCRAC experiment. Each bubble represents a separate day with key experimental steps, numbered based on the description in the protocol. The duration of each part is indicated in color-coded boxes: green for active hands-on time, and blue for mainly incubation hands-off steps. It is important to note that these durations are approximate and can vary depending on the experience and the quantity of samples being processed. The time duration given here should be approximately for 6 samples.

Like all UV-crosslinking-based methods, reCRAC is not without its limitations. The efficiency of crosslinking is expected to be variable, potentially displaying a bias for preferential binding of pyrimidines to specific amino acids ^13^. The efficiency of crosslinking can also fluctuate depending on the medium used, the cell type being studied, and the target protein itself ^6^. Additionally, the process of epitope tagging, essential for reCRAC method, can potentially influence protein stability, expression level, or RNA-binding efficiency. We therefore recommend confirmation of tagged protein synthesis and, if possible, function prior to reCRAC. Notably, reCRAC has currently been applied only in yeast cells and requires a significant quantity of starting material. However, the technique should be applicable to other organisms, with suitable adjustments to cell lysis conditions.

In our hands, reCRAC has demonstrated its efficacy in precisely mapping RNA binding sites for the yeast protein Pin4. This largely uncharacterized RNA-binding protein was previously implicated in the response to different stresses, particularly glucose withdrawal ^17^. In standard CRAC procedure, Pin4 suffered a high level of protein degradation (**Figure 2A, 2B**), which was greatly reduced by reCRAC (**Figure 2A, 2B**). We also observed improved cDNA library signal (**Figure 2C)** and improved signal-to-noise ratios (**Figure 2D**). We believe that reCRAC is a valuable tool for identification of biologically relevant protein-RNA interaction sites.

**Figure 2.**
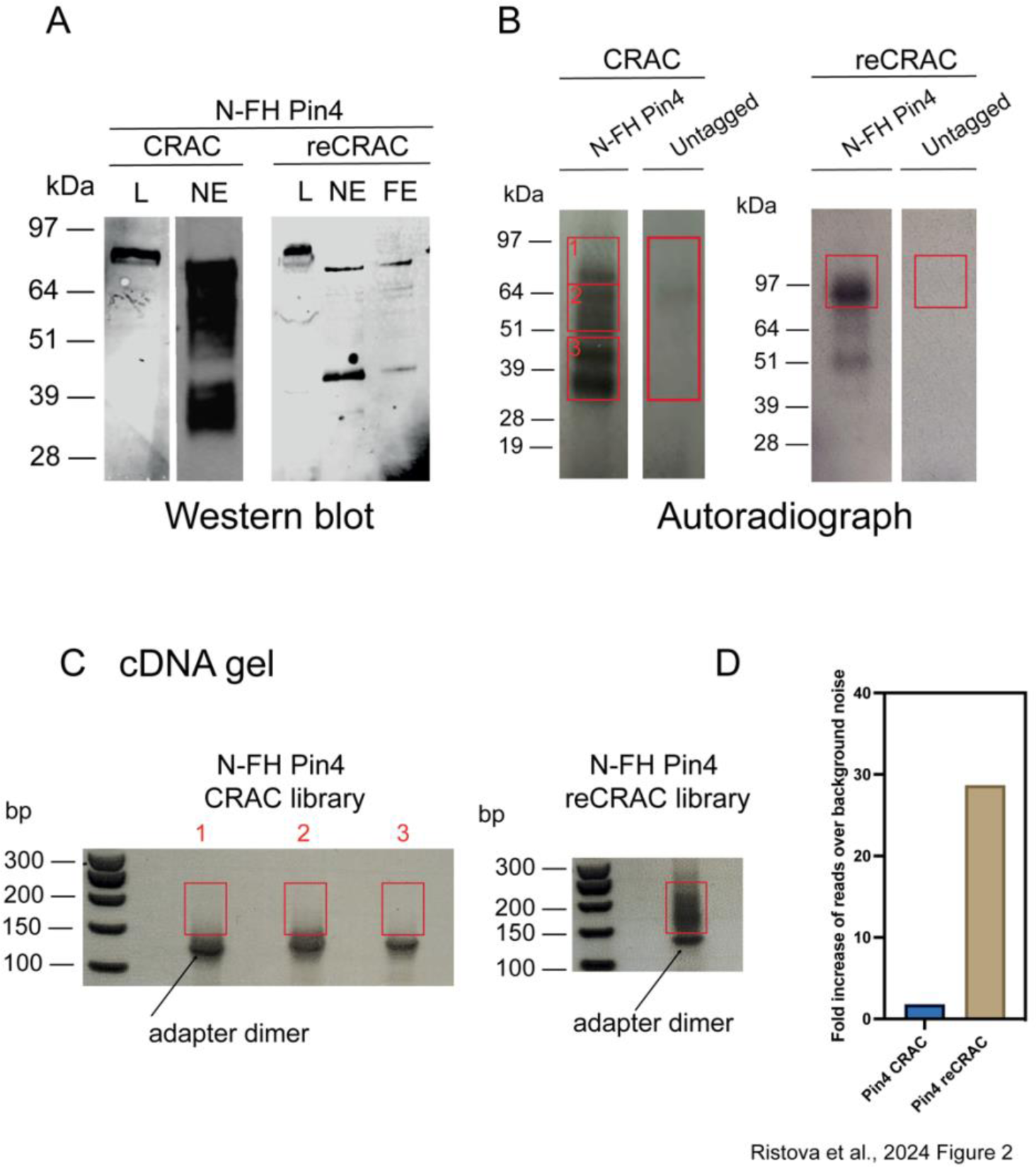
CRAC and reCRAC analysis of Pin4. A: Western blot analysis showing the N-FH (Flag-4xAla-8xHis) tagged Pin4 protein in the lysate (L) and after the elution from Nickel purification step (NE), and Flag elution step (FE) in CRAC and reCRAC. The protein is visibly degraded to a very large extent during CRAC, while in reCRAC, the degradation is reduced to two bands. B: Autoradiograph indicating the presence of covalently crosslinked N-FH Pin4 - RNA complexes in both CRAC and reCRAC. The red boxes indicate the regions excised for further steps. Three distinct regions (1, 2, 3) have been marked for excision in CRAC. The untagged strain was used as a negative control. C: Metaphore agarose gel electrophoresis of the resulting cDNA library from the CRAC and reCRAC method. The regions: 1, 2, 3; in CRAC cDNA gel correspond to the bands excised from the autoradiograph. The smear indicating the presence of cDNA products is readily visible in reCRAC and shows a good length distribution. Regions excised for sequencing are indicated in red. The lower strong band represents adapter dimer artifacts. E: Fold increase of reads over background (untagged strain) in CRAC (left bar) versus reCRAC (right bar).

## 2. Materials

In preparation for the experiment, a care should be taken to ensure that all materials, pipettes, and work surfaces are free of DNases and RNases. We recommend using RNase-free filter pipette tips throughout the experiment. The use of disposable gloves is mandatory throughout the process, and all laboratory practices should prioritize safety. When handling live cells, ensure a sterile environment to avoid contamination, and wash the bench and your gloves with 70% ethanol. All buffers should be prepared using DNase/RNase-free water. Once stock solutions are prepared, they should be filter-sterilized and stored at 4°C. Buffer A and Buffer B should be made fresh on the day of the experiment. Additives such as β-mercaptoethanol and protease inhibitors should be mixed into buffers immediately before use; these solutions are not suitable for overnight storage. Stock β-mercaptoethanol handling and phenol-chloroform extraction should be done in a fume hood. Follow all waste disposal regulations. Additionally, it is essential to carefully plan the experiment due to its length and labour-intensive nature, with some days requiring more than eight hours of laboratory work.

### 2.1. Yeast Strains and Culture Media

For the successful isolation of protein-RNA complexes, the protein of interest is tagged with either a N-terminal tag Flag-4xAla-8xHis (N-FH-protein), or a C-terminal tag 8xHis-4xAla-Flag (protein-HF-C). To explore the RNA partners of the uncharacterized *S. cerevisiae* RNA-binding protein Pin4, we engineered a yeast strain that expresses the N-FH tagged version of Pin4 from the endogenous *PIN4* gene locus. The tagged protein is therefore the only form of Pin4 in the cells, which showed no visible growth defects. The untagged parental yeast strain BY4741 was used as a negative control in our experiments.

Our yeast cultures were grown in a medium composed of 2% glucose (or other carbon source as required) and 2% yeast nitrogen base, supplemented with 2% amino acids excluding tryptophan (SD-Trp). The omission of tryptophan is critical as it absorbs light at 254 nm, the wavelength used for UVC crosslinking, potentially leading to reduced crosslinking efficiency.

### 2.2. Buffers and Solutions

1. DNase/RNase free water
2. Buffer A: 50 mM Tris-HCl pH 7.5, 300 mM NaCl, 0.1% NP-40, 4 mM imidazole pH 8.0, 6 M Guanidine-HCl, 5 mM β-mercaptoethanol
3. Digestion buffer: 50 mM Tris pH 7.8, 20 mM NaCl, 2 mM MgCl_2_, 5 mM β-mercaptoethanol
4. 1x PNK buffer: 50 mM Tris-HCl pH 7.8, 10 mM MgCl_2_, 0.5% NP-40, 5 mM β-mercaptoethanol. It can be stored at 4 °C long-term (several months) without β-mercaptoethanol
5. 5x PNK buffer: 250 mM Tris-HCl pH 7.8, 50 mM MgCl_2_, 25 mM β-mercaptoethanol
6. Elution buffer: 50 mM Tris-HCl pH 7.8, 500 mM NaCl, 0.3% SDS, 250 mM imidazole pH 8.0, 5 mM β-mercaptoethanol
7. TN150: 50 mM Tris-HCl pH 7.8, 150 mM NaCl, 0.1% NP-40, 5 mM β-mercaptoethanol. It can be stored at 4 °C long-term (several months) without β-mercaptoethanol
8. Buffer B: 50 mM Tris-HCl pH 7.8, 50 mM NaCl, 0.1% NP-40, 4 mM imidazole pH 8.0, 5 mM β-mercaptoethanol
9. Proteinase K buffer: 50 mM Tris-HCl pH 7.8, 50 mM NaCl, 0.1% NP-40, 4 mM imidazole pH 8.0, 5 mM β-mercaptoethanol, 1% SDS, 5 mM EDTA
10. 0.5 M EDTA-NaOH pH 8.0
11. 1 M Tris-HCl pH 7.5
12. 1 M Tris-HCl pH 7.8
13. 10% NP-40
14. 10% Triton X100
15. 100% and 70% ethanol
16. 10x TBE buffer: 890 mM Tris base, 890 mM boric acid, 20 mM EDTA
17. 14.3 M β-mercaptoethanol
18. 2.5 M imidazole-HCl pH 8.0
19. 20% SDS
20. 25:24:1 phenol-chloroform-isoamyl alcohol mixture
21. 3 M NaOAc pH 5.2
22. 5 M NaCl
23. Acetone
24. Guanidine HCl powder

### 2.3. Enzymes and Other Consumables

1. 100 mM ATP aliquots stored at -20°C
2. 100 mM DTT (Invitrogen, Cat#18080044)
3. 10x La Taq buffer (Takara, Cat#RR002M)
4. 2.5 mM dNTP mix (Takara, Cat#RR002M)
5. ^32^P-γATP (6000 Ci/mmol, Hartmann Analytic)
6. 5x first strand buffer (Invitrogen, Cat#18080044)
7. Benzonase (Merck, Cat#70664), Benzonase working stock prepared by diluting 25 U/µL of Benzonase in 1:250 of Digestion buffer with 50% glycerol, store long term at -20°C
8. DNA gel loading dye (6x), purple, no SDS (NEB, Cat#B7025S)
9. D (+)-Glucose Anhydrous (Thermo Scientific, Cat#10141520)
10. EDTA-free cOmplete protease inhibitor cocktail (Roche, Cat#11836170001)
11. Flag peptide (Sigma-Aldrich, Cat#F3290)
12. GlycoBlue (Invitrogen, Cat#AM9515)
13. GeneRuler 50 bp DNA ladder (Thermo Scientific, Cat#SM0371)
14. Kodak BioMax MS Autoradiography X-ray film (Cat#730-3224)
15. La Taq polymerase (Takara, Cat#RR002M)
16. Magnetic anti-Flag beads (Sigma-Aldrich, Cat#M8823)
17. Metaphore Agarose (Lonza, Cat#50180)
18. MinElute Gel extraction kit (Qiagen, Cat#28604)
19. MOPS running buffer 20x (Invitrogen, NP0001)
20. Ni-NTA Agarose beads (Qiagen, Cat#30410)
21. Nitrocellulose membrane (Thermo Scientific, Cat#88018)
22. NuPAGE bis-Tris 4-12% precast gradient gels (Invitrogen, Cat#NP0321BOX)
23. NuPAGE LDS Sample Buffer 4x (Invitrogen, Cat#NP0007)
24. NuPAGE transfer buffer 20x (Invitrogen, NP0006)
25. Proteinase K (Roche, Cat#3115836001)
26. Qubit dsDNA HS Assay Kit (Invitrogen, Cat#Q32851)
27. RNase H 5 U/µL (NEB, Cat#M0297S)
28. RNasin 40 U/µL (Promega, Cat#N2511, recombinant)
29. SeeBlue Plus2 Pre-stained Protein Standard (Invitrogen, Cat#LC5925)
30. Single dropout (SD)-Trp (Formedium, Cat#DCS0149)
31. Pierce Spin-columns (Thermo Scientific, Cat#69725)
32. SuperScript III (Invitrogen, Cat#18080044)
33. SYBR Safe (Invitrogen, Cat#S33102)
34. T4 PNK 10 U/µL (NEB, Cat#M02021S)
35. T4 RNA ligase I 10 U/µL (NEB, Cat#M0204S)
36. TSAP (Promega, Cat#M9910)
37. Yeast Nitrogen Base (Formedium, Cat#CYN0410)
38. Zirconia beads (Thistle Scientific, Cat#ZrOB05)

### 2.4. Specific Equipment

1. UV crosslinker. We are using the Vari-X linker ^18^. An alternative crosslinker such as the Megatron and Stratalinker device can be used ^15,19^.
2. Qubit 3.0 Fluorometer (Thermo Scientific).
3. Phosphoimager, able to print gel scan in its original size. We are using the Typhoon FLA9500 laser scanner (GE Healthcare Life Sciences).
4. Radiation room and authorization to work with radioactivity.
5. Geiger counter.

### 2.5. Oligonucleotides

Integrated DNA Technologies (IDT) supplied all oligonucleotides used in these experiments. Details of each are provided in **Table 1**. reCRAC requires Illumina-compatible 5’ and 3’ adapters and both RT (reverse transcription) and PCR forward and reverse primers. The 3’ adapter is a pre-adenylated miRCat-33 linker, the 5’ linker is barcoded and includes a sequence of three random nucleotides that helps in the identification of PCR duplicates and quantification during data analysis. The forward and reverse PCR primers contain specific sequences that are required for the binding of the amplified PCR products onto an Illumina flow cell.

**Table 1.**
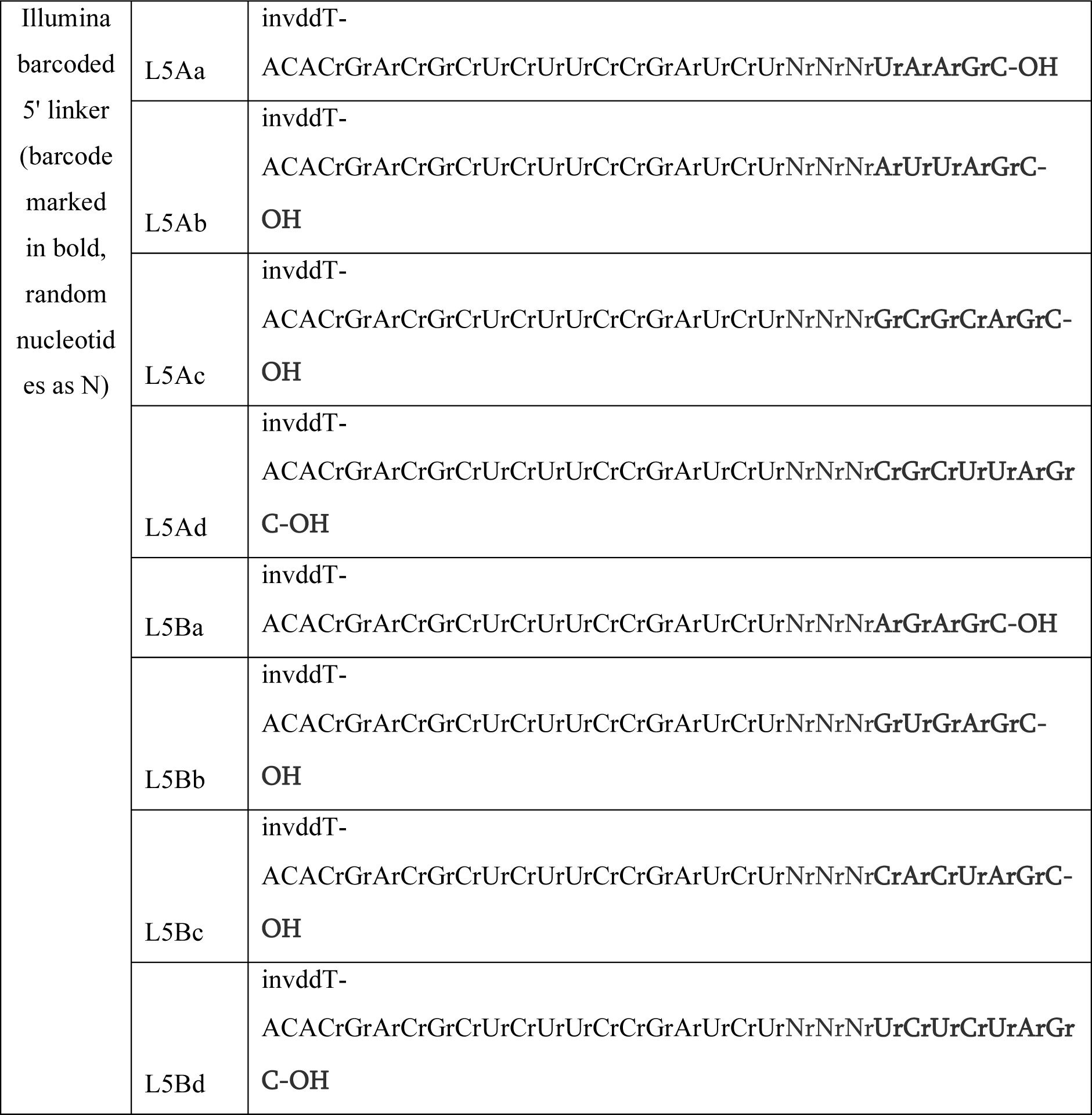
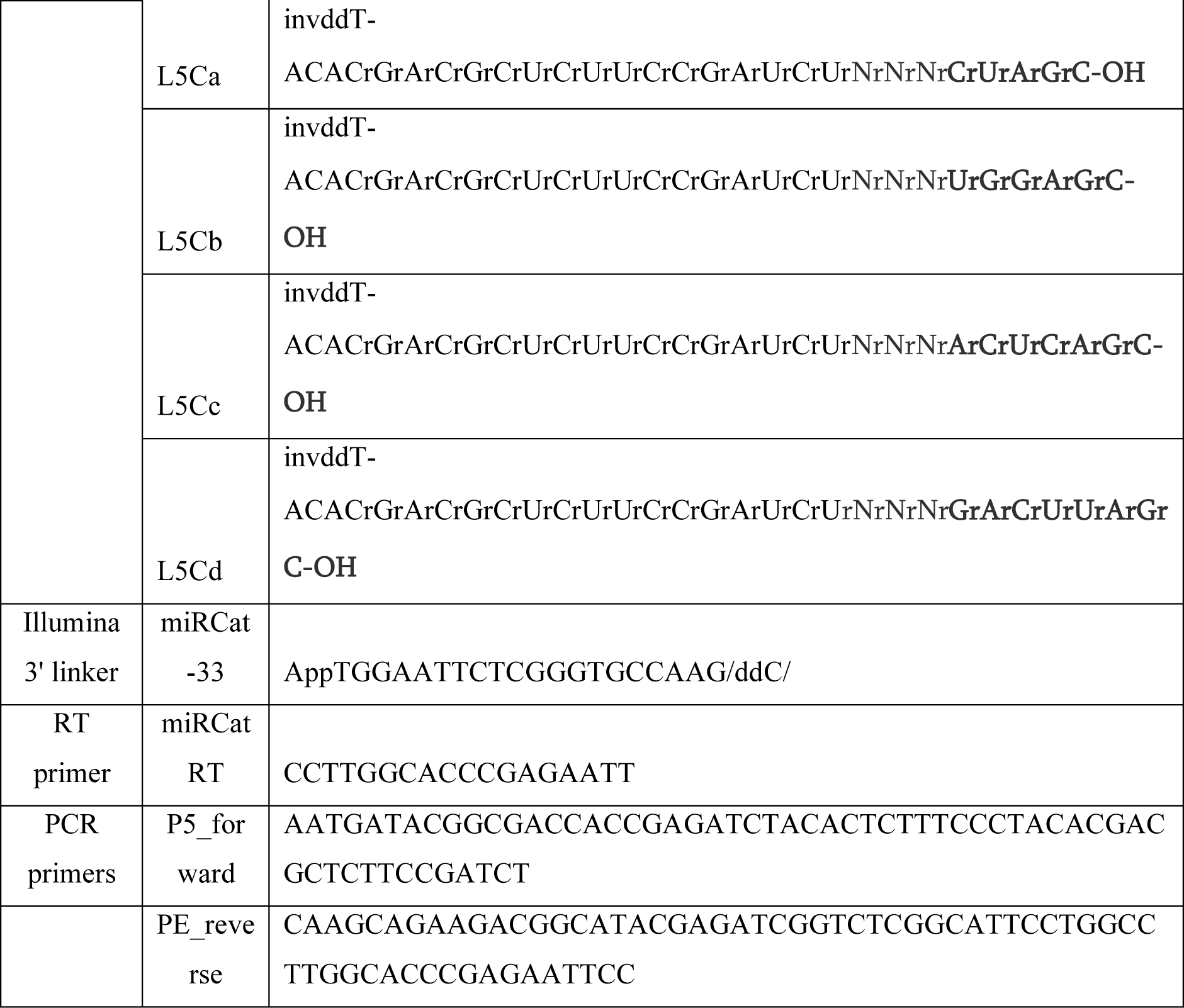
Oligonucleotides used in reCRAC.

For stability and to prevent degradation, these adapters are aliquoted and stored at -80°C.

## 3. Methods

### 3.1. Yeast Cell Culture and UV Crosslinking

1. Begin by streaking the FH-tagged yeast strains and the corresponding negative controls (e.g. untagged strain) from their glycerol stocks onto YPD agar plates (*see* **Note 1**). Incubate these at 30°C for approximately two days to allow colony growth.
2. Inoculate individual colonies into 70 mL of 2% single dropout (SD)-Trp medium, supplemented with 2% yeast nitrogen base and 2% glucose. Incubate the cultures overnight at 30°C with constant shaking at 200 rpm.
3. The following morning, dilute the overnight cultures to an optical density (OD_600_) of 0.05 in 800 mL of pre-warmed SD-Trp medium supplemented with 2% yeast nitrogen base and 2% glucose. Continue to incubate with shaking at 200 rpm until the culture reaches an OD_600_ of 0.45, which indicates a mid-log phase of growth. This depends on the strain and the sugar source; for BY4741, the time to reach the mid-log phase takes around 7 hours.
4. Prior to crosslinking, thoroughly cleanse the Vari-X crosslinker with water, and prewarm the UV lamps for 5 min to ensure optimal performance.
5. Crosslink using the Vari-X crosslinker set at 254 nm for 8 sec at room temperature with a dose of ∼150 mJ/cm^2^. This induces the formation of covalent bonds between proteins and the interacting RNA molecules.
6. Immediately following crosslinking, harvest the cells swiftly by filtration onto a nitrocellulose membrane. Put the filter into 50 mL of ice-cold water prepared in a 50 mL falcon tube and resuspend the cells by shaking the tube vigorously.
7. Carefully remove the membrane from the falcon tube, and centrifuge the cells down at 4,500 x *g* for 2 min at 4°C.
8. Remove the supernatant, collect any remaining water droplets with a vacuum pump, freeze the pellets and store them at -80°C.

### 3.2. Cell Lysis

The cell lysis process is conducted **on ice**, to prevent degradation of the cellular components.

1. Resuspend the frozen yeast cell pellets in 800 µL of Buffer A supplemented with 20 mM β-mercaptoethanol and a EDTA-free cOmplete protease-inhibitor cocktail (1 tablet per 50 mL of Buffer A).
2. Put 1.5 mL of Zirconia beads to the cell suspension.
3. Lyse the cells with six pulses of 1 min on a benchtop vortexer. Between each pulse, allow samples to cool **on ice** for 1 min to prevent heat build-up.
4. Add 2.4 mL of Buffer A **without** β-mercaptoethanol containing protease inhibitors. To completely mix the sample, vortex the tube vigorously for 10 sec using the benchtop vortexer, and then centrifuge at 4,500 x *g* for 5 min at 4°C to separate the lysate from the cell debris.
5. Transfer the supernatant carefully (∼ 4.5 mL) to three 1.5 mL locking cap tubes. Care is needed as the supernatant will be foamy due to the presence of guanidium. Spin the lysate for 20 min at maximum speed in a microcentrifuge at 4°C. Collect the three supernatants from each sample and combine to a single tube per sample, mix briefly.
6. Take 20 µL as an “Input” lysate for a western blot analysis (*see* **Note 2**). Guanidium cannot be loaded directly onto the gel, therefore, the sample is precipitated overnight. Dilute the input with 80 µL of TN150 buffer, then add 900 µL 100 % ethanol, and put to -20°C overnight. The following day, centrifuge the input at 13,000 x *g* for 10 min at 4°C. Wash the resulting pellet once with 70% ethanol to remove impurities, then resuspend in 1x LDS Sample buffer with 2% β-mercaptoethanol. Store at -20°C until ready to be run with Nickel and Flag eluates.

### 3.3. Nickel Bead Purification

All steps are carried out at room temperature unless specified otherwise.

1. Pre-equilibrate the Ni-NTA beads with Buffer A including 5 mM β-mercaptoethanol by washing the beads twice in 10 volumes of buffer; centrifuging at 400 x *g* to pellet the beads, and carefully discarding the supernatant after each wash. Make the pre-equilibrated Ni-NTA beads mixture by diluting the beads in 50:50 beads:buffer ratio.
2. Combine the lysate with 350 µL of the pre-equilibrated Ni-NTA beads mixture (containing 175 µL of beads) in a 5 mL Eppendorf tube. Use wide orifice pipette tips or cut the end of a tip with a sterile scalpel for more efficient pipetting of the beads.
3. Put the samples on the rotating wheel for 2 hours at room temperature.
4. Transfer the bead-lysate mixture into a Pierce spin-column. Let it stand for a minute or until the filter is visibly saturated and wet. Centrifuge the column for approximately 5 sec at a maximum of 300 x *g.* Following this speed is important as higher speed spins can damage the beads.
5. Wash 3x with 400 µL Buffer A including 5 mM β-mercaptoethanol with gravity flow. Spin down the final wash at 300 x *g* for 5 sec (*see* **Note 3**).
6. Wash 3x with 400 µL Digestion buffer. During the first wash, carefully rinse the inner walls of the column to remove leftover denaturing buffer. Spin down the columns after the final wash for 5 sec at 300 x *g*.
7. Seal the bottom of the column with the cap provided by the manufacturer.
8. Place the columns in a 37°C heat block.
9. Add 100 µL Digestion buffer, pre-warmed to 37°C and supplemented with 5 µL Benzonase working stock to the column.
10. Incubate the column for 5 min at 37°C with shaking at 900 rpm every 10 sec to ensure adequate mixing of beads with the digestion solution.
11. After nuclease treatment, open the lid carefully and remove the cap. The cap can be re-used for subsequent incubation steps with the same sample. Quickly centrifuge the column at 300 x *g* for 30 sec to remove the nuclease solution. Exercise caution to avoid contamination of gloves with the enzyme. Transfer the column to a new collection tube.
12. Immediately add 250 µL of Buffer A including 5 mM β-mercaptoethanol directly onto the beads at the bottom of the column to deactivate any residual nuclease.
13. Wash the column 2x with 400 µL of Buffer A including 5 mM β-mercaptoethanol.
14. Wash the beads 3x with 400 µL of cold 1x PNK buffer including 5 mM β-mercaptoethanol. From this point on, all steps should be carried out **on ice**. During the first wash, make sure to rinse the inner wall of the column to remove any traces of guanidium.

### 3.4. Alkaline Phosphatase Treatment of RNAs

1. Spin out the remaining buffer. Securely place a cap on the column and place into a clean Eppendorf tube. Prepare 80 µL of the following mix (set up at room temperature): 16 µL of 5x PNK buffer, 8 µL of TSAP, 2 µL of RNasin, and 54 µL of nuclease-free water (*see* **Note 4**).
2. Close the lid on the column and gently mix the contents by flicking the column with a finger. The beads should be visibly resuspended.
3. Place the column in a thermal mixer and incubate at 37°C for 30 min. Set the mixing to 800 rpm for 10 sec every 5 min.
4. Following the incubation, carefully open the column lid and remove the cap from the bottom. Wash the beads **on ice** once with 400 µL of Buffer A including 5 mM β-mercaptoethanol to inactivate TSAP.
5. Wash the beads 3x **on ice** with 400 µL of 1x PNK buffer including 5 mM β-mercaptoethanol to get rid of guanidium.

### 3.5. On-Bead Ligation of miRCat-33 DNA Linker to 3’ end

1. Spin out the remaining buffer, securely place a cap on the column and place into a clean 1.5 mL Eppendorf tube. Prepare ligation mix (80 µL final volume), adding T4 RNA ligase I last. The mix includes 16 µL of 5x PNK buffer, 8 µL of adenylated 3’ miRCat-33 linker, 2 µL of RNasin, 50 µL of nuclease-free water and 4 µL of T4 RNA ligase I (*see* **Note 5**).
2. Allow the ligation to proceed for 6 hours or overnight at 16°C (recommended). Set the mixer to agitate at 800 rpm for 10 sec every 5 min.
3. Following the incubation, open the column-lid with care and remove the bottom cap. Wash the beads **on ice** once with 400 µL of Buffer A including 5 mM β-mercaptoethanol.
4. Wash the beads **on ice** 3x with 400 µL of 1x PNK buffer including 5 mM β-mercaptoethanol to eliminate any traces of guanidium. Ensure that the first wash circulates around the rim of the column.

### 3.6. RNA 5’ end Phosphorylation and On-Bead Ligation of 5’ Linker

1. Spin out the remaining buffer, securely place a cap on the column, and transfer into a clean 1.5 mL tube. Add the following mix: 16 µL of 5x PNK buffer, 4 µL of T4 PNK, 3 µL of [^32^P]-γATP (10 μCi/μl), and 56 µL of nuclease-free water (*see* **Note 6**). We recommend preparing the mix without radioactive ATP, and then adding the radioisotope in a designated lab area for radioactive work.
2. Incubate the reaction for 40 min at 37°C with 800 rpm mixing every 5 min for 10 sec.
3. Add 1 µL of unlabelled 100 mM ATP for an additional 20 min to ensure thorough phosphorylation of the RNA 5’ ends.
4. At room temperature, wash the beads 3x with 400 µL of Buffer A with 5 mM β-mercaptoethanol. We recommend to carefully change the elution tube with each wash to minimize radioactive contamination.
5. Wash the beads with 3-4x with 1x PNK buffer containing 5 mM β-mercaptoethanol until radiation levels in the flow-through drop below ∼ 30 counts per second.
6. Spin out the remaining buffer and add 80 µL of ligation mix: 16 µL of 5x PNK buffer, 0.8 µL of 100 mM unlabelled ATP, 2 µL of **uniquely barcoded** 100 µM 5’ linker, 2 µL of RNasin, 4 µL T4 RNA ligase I, and 55.2 µL of nuclease-free water (*see* **Note 7**).
7. Incubate at 16°C for 4 hours or overnight (recommended) with mixing at 800 rpm for 10 sec every 5 min to facilitate the ligation of the 5’ linker to the RNA’s phosphorylated 5’ end.

### 3.7. Nickel Bead Elution

1. Wash the beads 3x with 400 µL of Buffer A including 5 mM β-mercaptoethanol.
2. After spinning out the void volume, perform the following elution twice using Elution buffer: for each elution, place a cap on the column, incubate the beads with 100 µL of Elution buffer for 10 min at room temperature with mixing at 750 rpm for 10 sec every minute. Spin into a new Eppendorf tube, pool eluates.
3. Take 10 µL of the Nickel eluate, add 10 µL of 2x LDS Sample buffer with 4% β-mercaptoethanol, boil for 5 min, and store at -20°C until ready to run on a gel. This is the “Nickel eluate” checkpoint for the western blot.
4. Dilute the eluate sixfold to make it compatible with Flag bead binding; use 50 mM Tris (pH 7.8) with 1% Triton X100, reducing SDS concentration to 0.05%, NaCl to 83.3 mM, and Imidazole to 41.7 mM.

### 3.8. Flag Bead Purification

Maintain **a cold environment** throughout this procedure.

1. Prepare 100 µL of magnetic anti-Flag beads suspension per sample (containing 50% of beads) pre-washed twice with 5 mL of TN150 buffer. The washes are performed by first collecting the beads with a brief pulse spin to 1,000 rpm in a microcentrifuge, and then using a magnetic rack to remove supernatant. After the final wash, resuspend 50 µL of the beads in TN150 buffer to 100 µL final.
2. Combine the diluted eluate from step 4 of **section 3.7**. with the pre-washed beads and place on a rotating wheel for 1.5 hours at 4°C to bind (*see* **Note 8**).
3. After the incubation, pellet the beads by spinning them up to 1,000 rpm in a microcentrifuge. Use a magnetic rack to remove the supernatant.
4. Resuspend the beads in 1 mL of TN150 buffer and transfer them to a new 1.5 mL tube. Rotate for 5 min at 4°C for uniform washing.
5. Repeat the wash twice with 1 mL of TN150.
6. Elute twice with 150 µL of Flag peptide, prepared at 100 µg/mL in 1x TN150. For each elution, incubate the beads at 37°C for 15 min with vigorous shaking at 1,200 rpm.
7. Combine the eluates post-elution and immediately cool **on ice** for 1 min.
8. For the western blot, take 10 µL of the Flag eluate, add 2x LDS Sample buffer with 4% β-mercaptoethanol, boil for 5 min, and store until ready to run on a gel. This is the “Flag eluate” checkpoint for the western blot. When the western blot analysis of Input, Nickel, and Flag eluates is ready to be run, heat the samples at 95°C for 5 min, briefly centrifuge. Load the samples onto a 1.5 mm thick, 10 well NuPAGE 4-12% gradient gel. Run the gel with 1x NuPAGE MOPS running buffer with SeeBlue2 pre-stained protein standard gel for 1 hour at 150V. Transfer onto a nitrocellulose membrane using the 1x NuPAGE transfer buffer with the parameters optimized to the protein of interest. Block the membrane, perform the primary and secondary antibody incubation, and visualise the membrane.

### 3.9. Gel Electrophoresis

1. Add 2 µL GlycoBlue, 4 volumes of acetone (1.2 mL) to the combined eluate from step 7 of **section 3.8**., and incubate for at least 2 hours or overnight, at -20°C.
2. Centrifuge the samples at max speed in a microcentrifuge for 20 min at 4°C. Carefully remove the supernatant ensuring that the small blue pellet remains undisturbed. No wash of the pellet is required, unless the pellet appears large and white, in which case an additional acetone wash is needed. Allow the pellets to dry at room temperature with the lids open for a few minutes. Avoid over-drying, which can impede resuspension.
3. Gently resuspend the pellet in 30 µL 1x LDS Sample buffer including 2% β-mercaptoethanol. Pipette up and down, checking resuspension progress with a Geiger counter (*see* **Note 9**).
4. Heat the samples at 65°C for 10 min. Briefly centrifuge post-heating.
5. Load the samples onto a 1.5 mm thick, 10 well NuPAGE 4-12% gradient gel, using 1x NuPAGE MOPS running buffer. Add a protein ladder such as SeeBlue2 pre-stained protein standard.
6. Electrophorese at 150V for 1 to 1.5 hours, or until the dye reaches the foot of the gel.
7. After the run is finished, dismantle the gel cassette, retaining the gel on one of the plastic sides (*see* **Note 10**). Wrap in a saran wrap, place in an X-ray film cassette, secure with tape (crucial to avoid movement!) and expose to X-ray film for 1 hour or overnight at - 80°C, depending on the radioactive signal intensity. Include a chemiluminescent marker to aid in subsequent alignment of the film with the gel. The most sensitive film, MS film should normally be sufficient for detection using a 1-hour exposure. Overnight X-ray film exposure might be necessary.
8. Develop the X-ray film and align it with the gel using the chemiluminescent markers as guides. Excise the region corresponding to the size of your target protein plus an additional 20 kDa, to account for RNA crosslinked to the protein of interest. Excise the equivalent region of the negative control lane. Put the excised fragments of the gel into a clean 1.5 mL tube (*see* **Note 11**).
9. Smash the excised gel sections into sub-millimetre pieces with a bent P200 pipette tip exercising caution to prevent gel fragments from scattering out of the tube. Crush the pieces against the tube walls with the tip until they have a paste-like consistency. Bending the tip beforehand helps to avoid accidental suction of gel pieces during this process.

### 3.10. Proteinase K Treatment

These steps are crucial to release RNA from protein-RNA complexes.

1. Incubate the gel slices in 600 µL of Buffer B including 1% SDS and 5 mM EDTA. Add 100 µg (5.0 µL) Proteinase K from a frozen stock solution of 20 mg/mL in water and incubate at 55°C for 2 hours with vigorous shaking at 1,100 rpm to ensure thorough enzymatic digestion.
2. To remove the gel fragments, transfer the digested mixture to a Pierce spin-column. If necessary, cut the end of a 1 mL pipette tip with a sterile scalpel to make the transfer of the gel chunks easier. Centrifuge the column at the maximum speed for 1 min. Check to ensure all the liquid passes through, leaving gel pieces trapped on the filter.
3. Carefully transfer the supernatant to a new tube. Add 75 µL of 3 M NaOAc pH 5.2, and 750 µL of phenol:chloroform:isoamyl alcohol (24:25:1) in the fume hood.
4. Vigorously vortex the mixture, then centrifuge at room temperature for 20 min at maximum speed. Carefully collect the aqueous top layer to a new tube, taking care to avoid the bottom layer or the interphase. Repeat the phase separation process using the same volumes.
5. Precipitate the RNA with 1.5 mL ice-cold 100% ethanol and 2 µL of GlycoBlue. Incubate the samples at -80°C for 30 min or at -20°C overnight.
6. Centrifuge the samples at 4°C for 20 min at maximum speed in a microcentrifuge. Wash the RNA pellet with 750 µL of 70% ethanol, vortex, and centrifuge again for 5 min at 4°C at maximum speed. Allow the pellet to air-dry for ∼5 min.

### 3.11. Reverse Transcription of Purified RNAs

This will generate cDNA from the extracted RNAs.

1. Resuspend the RNA pellet in the following mix: 4 µL of 2.5 mM dNTP mix, 1 µL of 10 µM RT primer, and 8 µL of nuclease-free water.
2. Heat this mixture to 80°C for 3 min. Immediately after, snap chill the samples **on ice** for 5 min (*see* **Note 12**).
3. Collect the contents by brief centrifugation and add 6 µL of the following mix: 4 µL of 5x first strand buffer, 1 µL of 100 mM DTT, and 1 µL of RNasin.
4. Incubate the mixture at 50°C for 3 min to allow the RT primer to bind to the RNA. Subsequently add 1 µL of SuperScript III reverse transcriptase. Incubate the reaction for 1 hour at 50°C, which will synthesize the cDNA from the RNA template.
5. Following the reverse transcription, inactivate the SuperScript III enzyme by heating the sample to 65°C for 15 min.
6. To remove the RNA template, add 2 µL of RNase H and incubate at 37°C for 30 min.

### 3.12. PCR Amplification of cDNA

1. Set up three PCR reactions per sample (*see* **Note 13**), combining the following: 5 µL of 10x La Taq buffer, 1 µL of 10 µM P5_forward primer and 1 µL of 10 µM PE_reverse primer, 5 µL of 2.5 mM dNTPs, 0.5 µL of La Taq, 3 µL of reverse transcription reaction, and 34.5 µL of nuclease-free water.
2. Program the PCR thermocycler as follows:

3. Pool and precipitate the three PCR products per sample: **on ice** add 1 µL of GlycoBlue, 0.1 volume of 3 M NaOAc, and 2.5 volumes of 100% ice-cold ethanol.
4. Put the mixture to -80°C for 30 min, then spin for 20 min at 4°C.
5. Wash the pellet with 70% ethanol, spin for 20 min at 4°C, and air-dry for 5 min.
6. Resuspend the pellet in 15 µL of nuclease-free water.

|  |
| --- |
| 95°C for 2 min |
| <u>20-25 cycles of:</u> |
| 98°C for 20s |
| 52°C for 20s |
| 68°C for 20s |
| 72°C for 5 min |

### 3.13. DNA Gel Electrophoresis and Purification

1. Add 3 µL of 6x DNA loading dye to the sample.
2. Prepare a 3% Metaphore agarose gel by first soaking agarose in 1x TBE for 30 min. Heat it in a microwave, until all of the agarose is dissolved. Once it is cooled to approximately body temperature, add 1:10,000 SYBR safe DNA stain for visualisation, and pour the gel. Allow the gel to solidify at room temperature, and then put it to 4°C for at least 30 min.
3. Load the entire volume of the samples alongside a 50 bp DNA ladder onto the prepared 3% Metaphore agarose gel. Run the gel on a metal block set in a container filled with ice to maintain a low temperature. Run the gel at 80 V until the dye front has migrated to the end of the gel. This usually requires ∼ 2 hours.
4. Scan the gel with a phosphoimager. The strong band at 120 bp are amplified linker-linker dimers, with the cDNA libraries forming a smear above this. Print the scanned image at 100% size.
5. Overlay the gel onto the printed scanned image placed below a glass slide. With a sterile scalpel, excise the section of the gel containing the cDNA library, typically running from 150 to 250 bp, while avoiding the 120 bp primer dimers (*see* **Note 14**).
6. Place the excised gel slices into a clean tube. Re-scan the gel to confirm that the intended region has been accurately removed.
7. Add 6 volumes of Buffer QG from the MinElute Gel Extraction Kit to 1 volume gel (100 mg ∼ 100 µL). Melt at 42°C for 10-15 min or until the gel slice has completely dissolved.
8. Transfer the liquid gel mixture to a MinElute column placed in a provided 2 mL collection tube and centrifuge for 1 min at 16,000 x *g*. Discard the flow-through and repeat the process with any remaining sample to ensure complete binding to the column.
9. Perform additional wash with 750 µL of Buffer QG. Spin for 1 min at 16,000 x *g*, and then discard the flow-through.
10. To remove any remaining impurities, add 750 µL of Buffer PE from the MinElute Gel Extraction Kit and let it sit for 10 min at room temperature. Optionally, you can invert the column to ensure all of Buffer QG will be washed away. Centrifuge again for 1 min at 16,000 x *g* and discard the flow-through.
11. Spin the empty columns at 16,000 x *g* for 1 min.
12. Transfer the columns to new, clean 1.5 mL tubes for elution.
13. Elute the DNA by adding 10 µL of pre-warmed (65°C) nuclease-free water to the column. Allow the water to penetrate the column matrix by letting it stand for a few minutes, then centrifuge for 1 min at 16,000 x *g*. Repeat the elution step.
14. Use a Qubit dsDNA HS Assay Kit to quantify the cDNA library. Store the purified libraries at -20°C until they are ready for sequencing.

### 3.14. Sequencing and Analysis

Libraries can be pooled and sequenced in a single run. We used an Illumina MiniSeq system, with single end sequencing and 75 bp read length, using the High Output Reagent Cartridge and followed the manufacturer’s instructions. Paired-end sequencing can be also performed, however, modifications to the bioinformatics analysis would be necessary.

Analysis of Pin4 sequencing data obtained from reCRAC was performed using custom scripts and the pyCRAC software packages ^16,20^. Below, we describe the main steps in processing of raw reads and the most commonly used pyCRAC tools. The analysis can be adapted for specific proteins and to address specific biological questions. The analysis was performed on a server using a Bash terminal.

#### 3.14.1. Debarcoding, Filtering and Adapter removal

The sequencing analysis starts by separating the reads originating from different samples/libraries into independent files, i.e. demultiplexing the raw output file, Output.fastq, by the 5’ linker barcodes, using the pyBarcodeFilter.py tool:

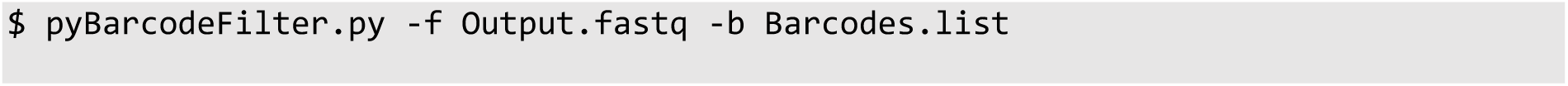

The Barcodes.list is a tab-delimited file that contains the list of barcodes and the names of the samples corresponding to the barcodes. Here is an example of the Barcodes.list used in reCRAC:

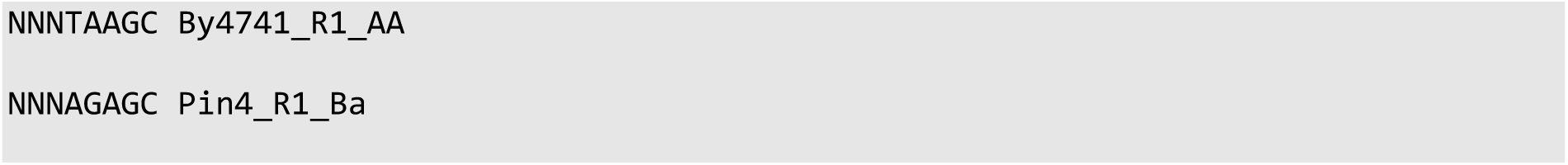

Where By4741_R1_Aa is the untagged control, and Pin4_R1_Ba is the target protein. The pyBarcodeFilter.py will pool all the reads with the same 5’ linker barcode into the same file, named according to the Barcodes.list. In addition, the random nucleotides present in 5’ linker are trimmed at this step and incorporated into the headers of each sequence within the resulting fastq file, which will be crucial during the subsequent Collapsing step. These random nucleotides help distinguish PCR duplicates from independent ligation events. Header example of the resulting fastq file from pyBarcodeFilter.py:

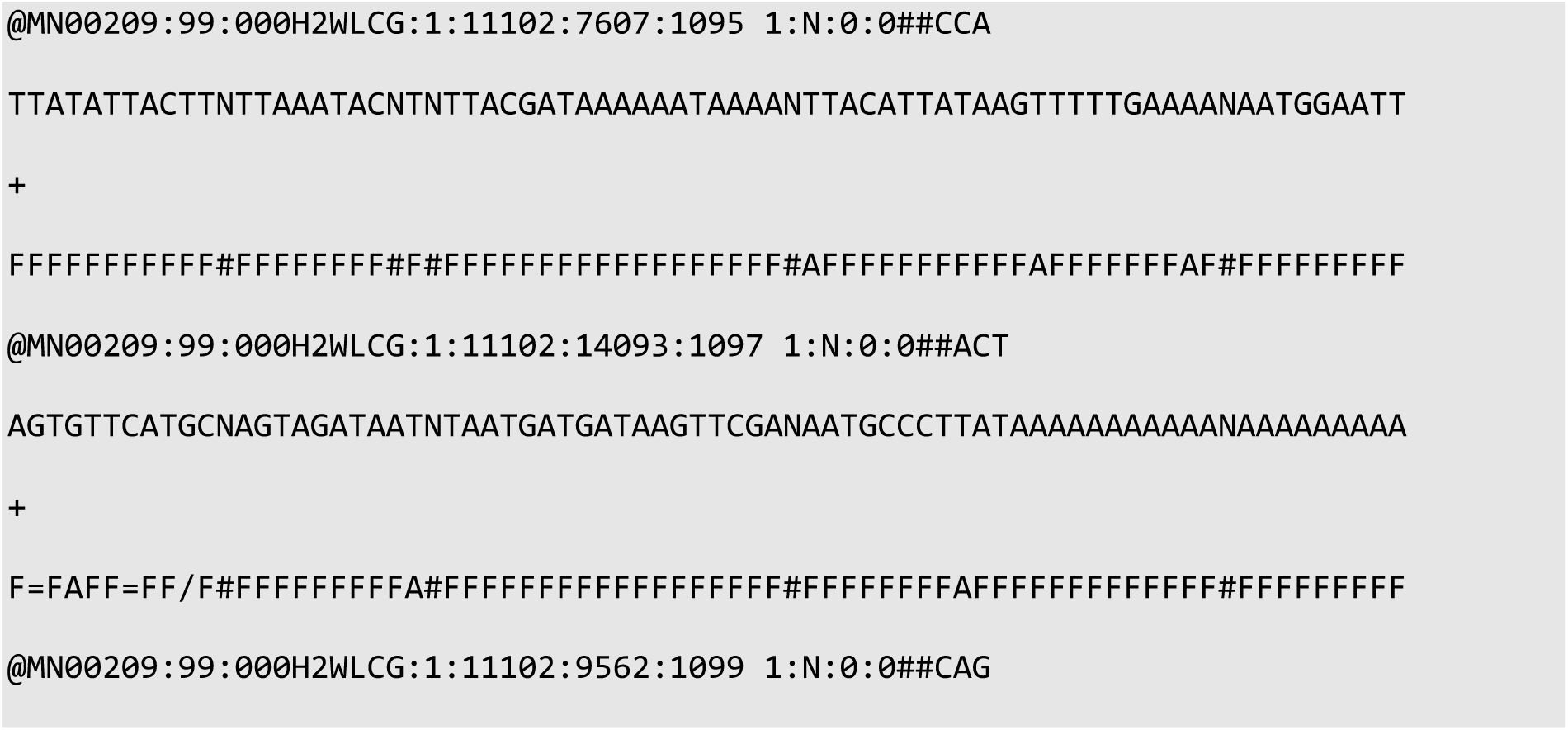

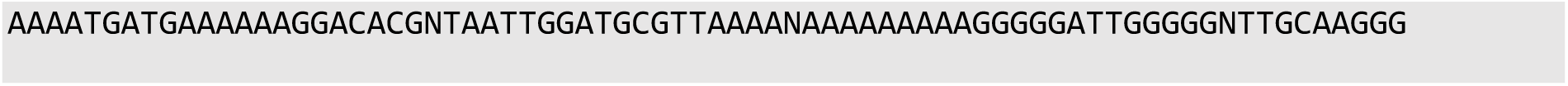

Quality of the sequencing data can be checked with FastQC:

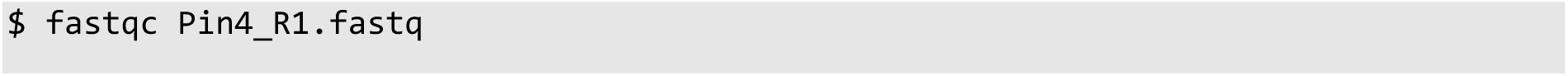

Alternatively, FastQC can be run on all fastq files in parallel:

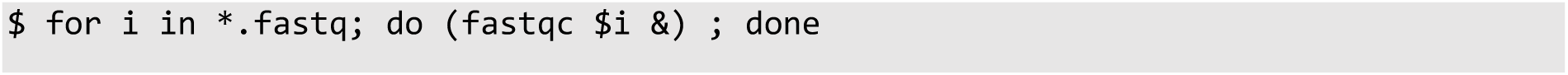

Next, the Flexbar tool is used for the quality filtering and adapter trimming ^21^:

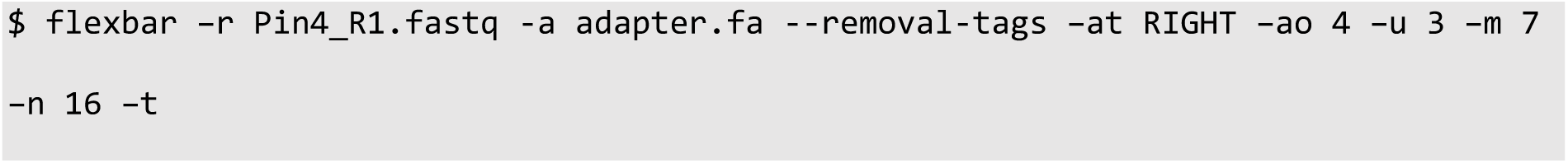

The file, adapter.fa, contains the sequence of the adapter to be trimmed (TGGAATTCTCGGGTGCCAAGGC). This is the sequence that was added on the 3’ end during cloning procedure and PCR amplification. The parameters are set to specify trimming process details. We advise performing a quality control with FastQC on the output files to confirm adapter removal and ensure that the fastq files passed the quality controls.

#### 3.14.2. Collapsing

This code removes reads that are predicted to be PCR duplicates, by collapsing reads with identical sequences including the three random nucleotides present in the 5’ linker, using the pyFastqDuplicateRemover.py tool. The following command iterates through all fastq files in the working directory and puts the output of each as collapsed.fasta file:

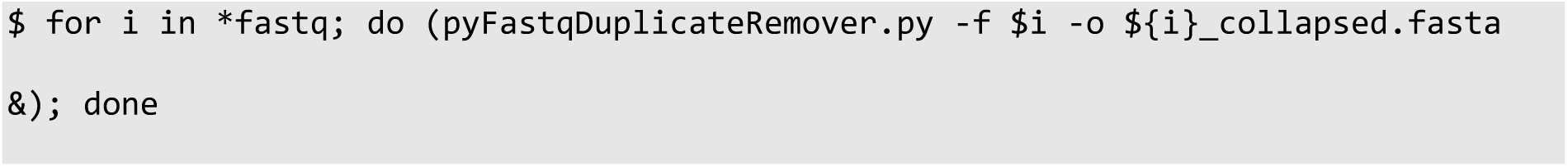

The example lines of the resulting fasta file from pyFastqDuplicateRemover.py:

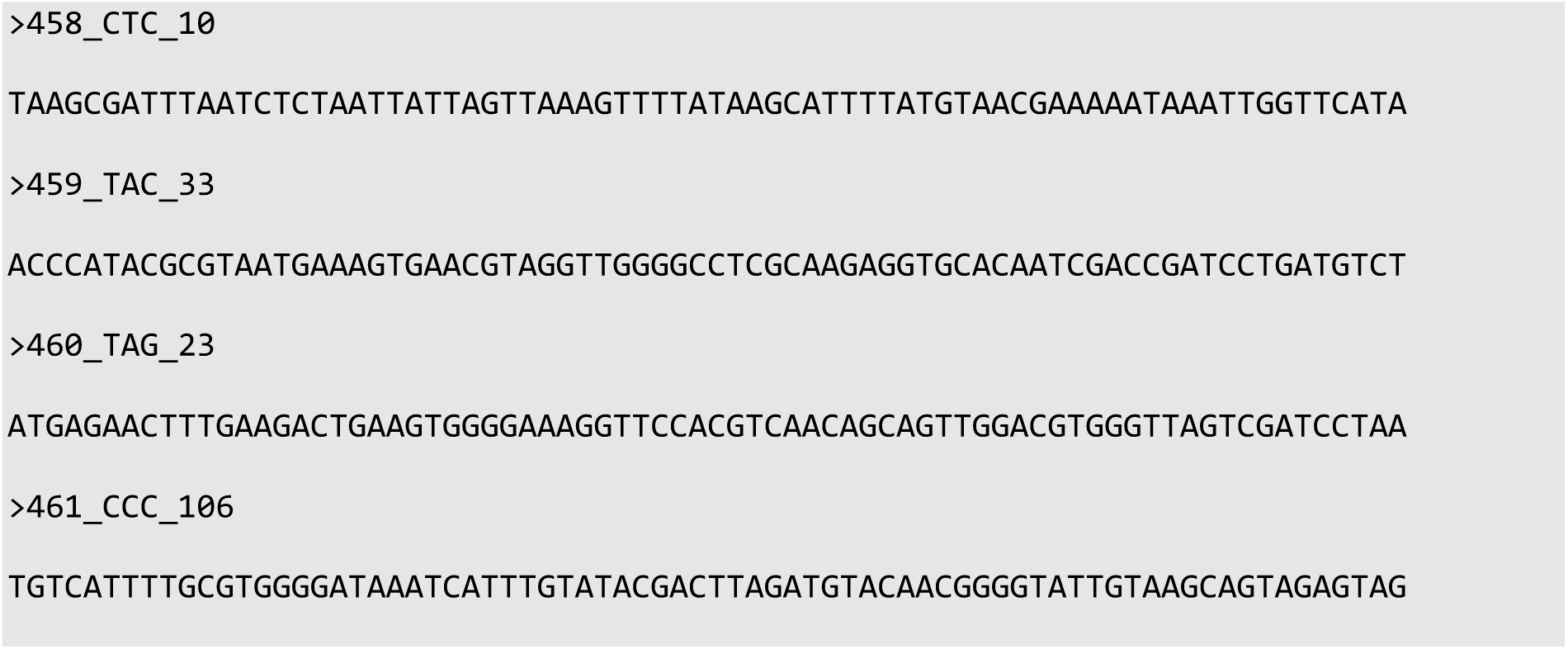

#### 3.14.3. Alignment

The collapsed reads are aligned to the reference genome of *Saccharomyces cerevisiae* (SGD v64) using the Novoalign tool (V2.07.00, Novocraft) with genome annotation from Ensembl (EF4.74) ^22^:

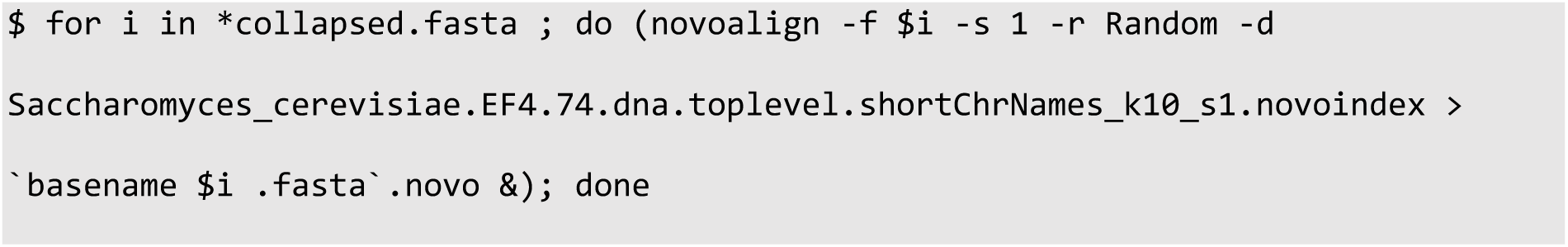

The Saccharomyces_cerevisiae.EF4.74.dna.toplevel.shortChrNames_k10_s1.novoindex is the genome-specific index file generated by novoindex with parameters -k10 s-1, fasta.novo is the ouput file. The resulting novo file has the following header and structure:

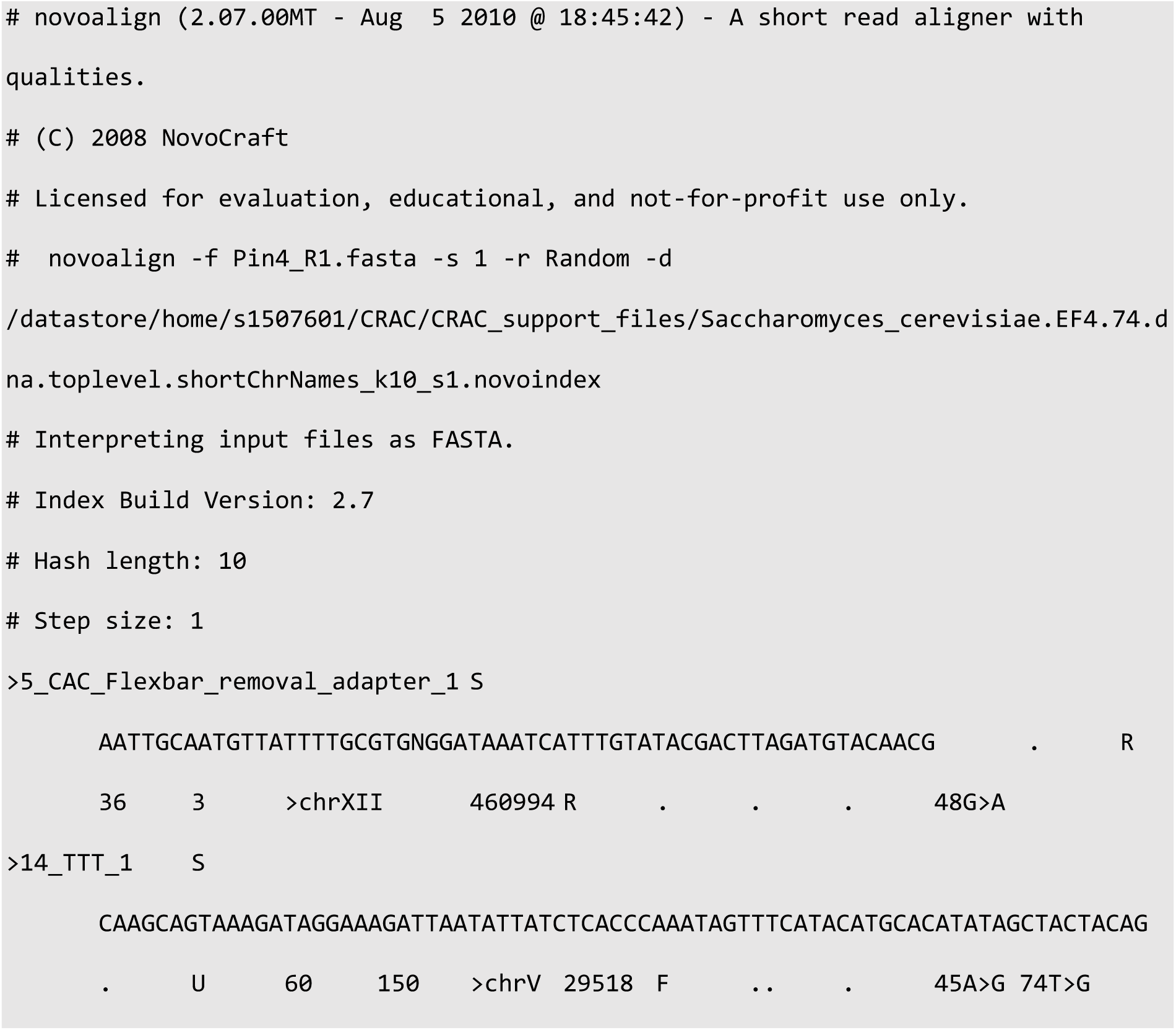

#### 3.14.4. Read Counting

All reads that align to each feature in the genome, such as introns, exons, CDS, are summed using the pyReadCounters.py from the pyCRAC package. It accepts input files in either .novo or .sam format, along with a gene transfer format (GTF) file that contains gene ID coordinates. For our analysis, we utilized the Ensembl GTF version EF4.74, which includes the mRNA annotations with UTRs, as well as different non-coding RNA categories such as CUTs and SUTs. Additionally, the pyReadCounters.py offers various options, including the type of normalization, read orientation, and alignment quality filtering. In the code below, `-- rpkm` option was used to RPKM normalize the reads:

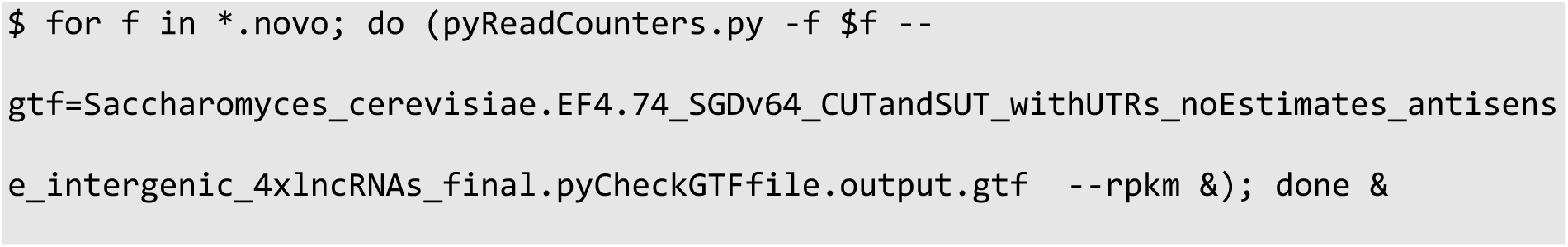

The script provides several outputs, including a read count file, with the RPKM normalized cDNA counts mapped to genomic features; GTF file containing additional information about the reads overlapping genomic features in a sample; and read statistics file providing the information about the complexity of the dataset.

#### 3.14.5. Data Visualization

We can visualise the protein binding sites across the whole genome using the Integrative Genomics Viewer (IGV) ^23^. Reads are aligned again using Novoalign with modified parameters to obtain the output in a different format (SAM):

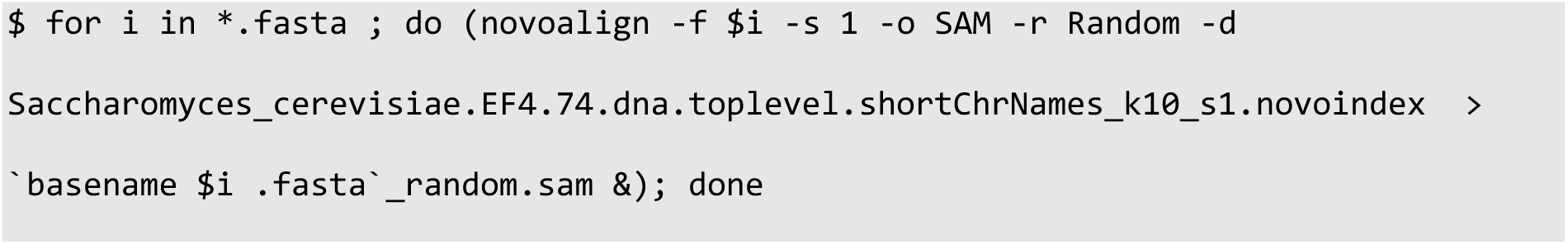

The SAM files are then converted into binary BAM files, which we sorted and indexed with ‘samtools’:

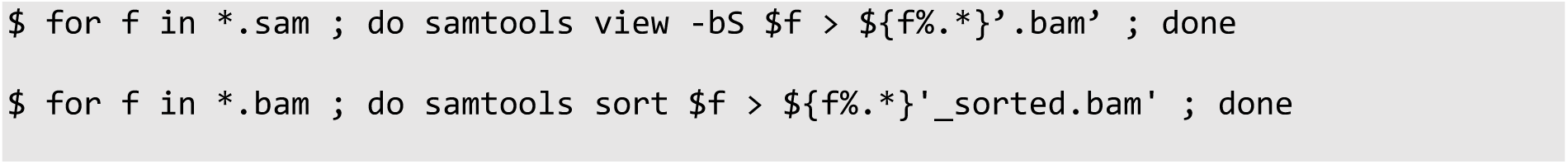

To prepare the data for visualization, a scaling factor is calculated to normalize the reads to the size of the library. This normalization factor is based on the number of mapped reads, often normalized to a standard number such as one billion reads. To find out how many reads were mapped:

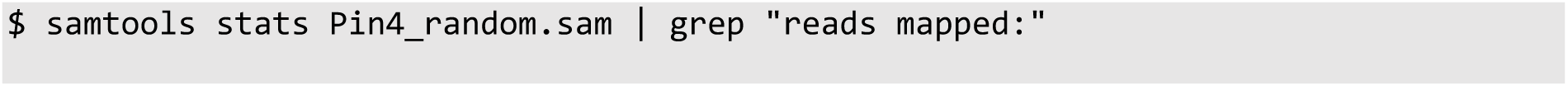

As an example of scaling factor calculation: Pin4_random.sam has 1,448,215 reads, dividing one billion by this number gives the scaling factor of 690.5. Using ‘bedtools’, coverage is computed separately for each strand and scaled by the calculated factor, generating bedgraph files:

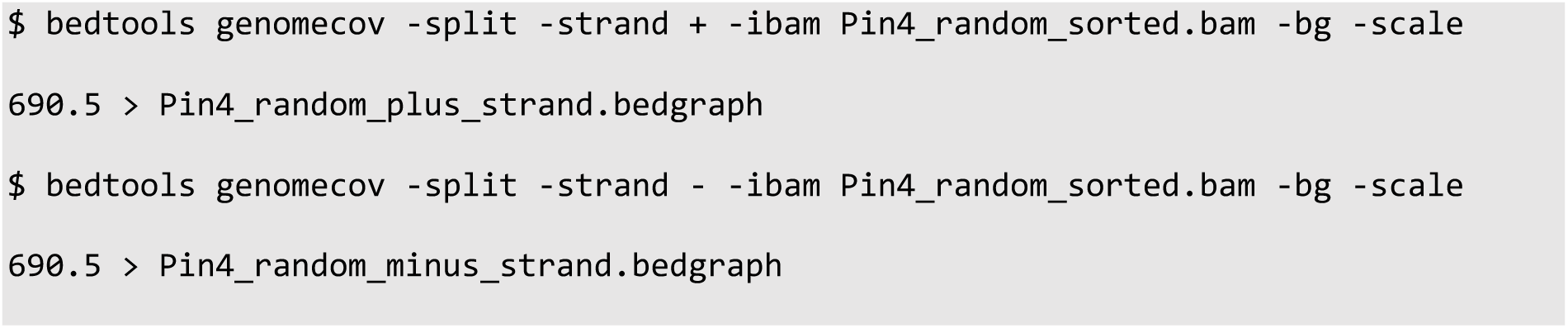

The bedgraphs files are then loaded into IGV.

Another useful data visualisation step involves using ’pyPileup.py’ tool to analyse the distribution of reads along specific genomic regions of interest. The pyPileup.py provides insights into the specific sites where crosslinks occur based on the number of deletions and substitutions at a nucleotide resolution. These result from nucleotide misincorporation or skipping during reverse transcription, at the site of the amino acid-nucleotide crosslink. The code needs the following inputs: a GTF output from pyReadCounters.py, and a text file containing the gene name(s) of interest. The code also contains the GTF annotation file with the standard genomic feature annotations, and the .tab file with the genomic reference sequence in a more concise format:

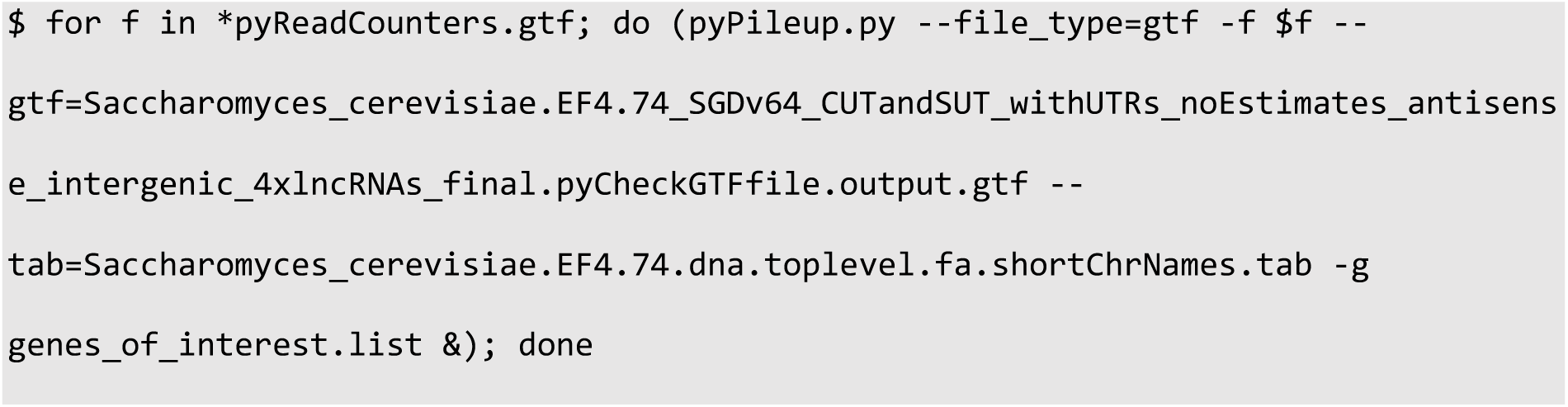

The output of this code is a tab-delimited file containing the number of deletions and substitutions per each nucleotide of the target gene.

### 4.0. Notes

1. It is advisable to incorporate a control sample within the experimental design in addition to experimental replicates. Suitable controls may include an untagged strain or a no-UV control. These controls are anticipated to yield negligible signals during SDS-PAGE analysis, and any libraries generated from sequencing should display significantly fewer reads than target samples; yielding at least 100-fold fewer unique cDNAs.
2. We collect samples from both the lysate and the eluate at each step of purification for western blot analysis. This approach allows monitoring of the target protein throughout the experiment and identifies the specific step should any problem arise.
3. For wash steps, allow the solution to pass through the column by gravity for a minimum 30 sec, before spinning them for approximately 5 sec at no more than 300 x *g* to avoid damaging the beads. Sometimes the columns exhibit slow or no flow by gravity, which we observed with certain batches. In this case, centrifuge the columns at 300 x *g* until all the liquid passes through.
4. Treating the RNAs with alkaline phosphatase will remove any 5’ phosphates left behind by the Benzonase cleavage of RNA to prevent self-ligation of RNA molecules.
5. The ligation utilizes a specialized 3’ miRCat-33 linker, which is chemically blocked at its own 3’ end (ddC) to prevent self-ligation, and pre-activated by adenylation at the 5’ end (App) to enable the ligation reaction without the addition of ATP.
6. Phosphorylation using radiolabelled ATP is initially employed to facilitate the detection of protein bound RNA on the SDS-page gel in the subsequent steps. This is followed by the addition of high concentration of non-radioactive ATP, allowing for efficient phosphorylation at all RNA 5’ ends, which is required for subsequent ligation to the 5’ linker.
7. The 5’ linkers are designed with a blocked 5’ end (invddT) to prevent self-ligation and incorporate unique barcodes for accurate sample identification, allowing for pooling multiple samples into one sequencing run in later step (**section 3.9**., step 3), and the identification of each original sample during the demultiplexing phase. It is crucial to use different barcodes for each sample. The 5’ linkers also include random nucleotides, enabling removal of PCR duplicates.
8. The samples will be still very radioactive at this point. We therefore recommend putting the rotating wheel into a fridge located in the radioactive lab if possible.
9. When analysing identical proteins under different conditions or across multiple replicates, this is the step when multiple samples can be pooled together. Begin by resuspending one sample pellet in 30 µL of 1x LDS Sample buffer, and then transfer everything to the second sample pellet, which will be merged during resuspension. Combining the samples at this step can minimize background noise and technical variations, ensuring consistent handling and uniform gel extraction in later steps.
10. Accurate alignment of the X-ray film over the gel is essential for precise excision of protein-RNA complexes. To ensure proper alignment, firmly secure the gel within the cassette using tape to prevent it from moving. Aligning the film to the upper left or right corner of the gel also aids in achieving an accurate cut.
11. An alternative method is cutting out from a membrane. This involves transferring the protein-RNA complexes from a gel to a nitrocellulose membrane, exposing the membrane to an X-ray film, and excising the band(s) from the membrane. This offers a cleaner result as documented by Delan Forino et al., 2019 ^16^. However, this approach comes with a risk of losing a large proportion of the protein-RNA complexes if transfer is inefficient.
12. Heating at 80°C will denature any secondary structures within the RNA, while snap chilling the samples on ice immediately after locks the RNA strands in a linear form, facilitating better primer annealing.
13. PCR reactions should be prepared in a UV-sterilized environment to minimize the risk of contamination. The goal is to achieve a total cDNA yield of 5 to 50 ng, which necessitates adjusting the number of PCR cycles accordingly. Typically, 21 cycles are adequate to produce diverse libraries from cDNA synthesized from Pin4-associated RNAs. If the cDNA is of low abundance, it may be necessary to increase the cycle count to approximately 23-25. However, it is essential to prevent overamplification, which can skew the representation of the library.
14. At the stage of DNA Gel Electrophoresis, adjusting the size distribution of the DNA library is possible. The size selection should be guided by the intended sequencing length, the nature of the protein under study, and the specific biological questions being addressed by reCRAC. If sequencing will be limited to e.g 80 base pairs, it is practical to exclude overly long cDNAs as they may reduce the resolution of protein-binding site identification. Conversely, it is generally beneficial to limit the presence of very short sequences (under 20 nucleotides) in the library since they can be challenging to map with confidence, and they are preferentially bound by the flow cell, introducing a bias.

## Acknowledgements

We thank Aleksandra Helwak for critical reading of the MS. MR was supported by an EASTBIO PhD fellowship [543KOM/G40389]. VS and DT were supported by Wellcome Principal Research Fellowships [109916, 222516]. This work was facilitated by funding for the Wellcome Discovery Research Platform for Hidden Cell Biology [226791] and we gratefully acknowledge support from the Microscopy and Bioinformatics cores. Work in the Wellcome Centre for Cell Biology was supported by Centre Core Grants (092076 and 203149).

## References

1. Beckmann, B.M., Horos, R., Fischer, B., Castello, A., Eichelbaum, K., Alleaume, A.-M., Schwarzl, T., Curk, T., Foehr, S., Huber, W., et al. (2015). The RNA-binding proteomes from yeast to man harbour conserved enigmRBPs. Nat. Commun. 6, 10127. 10.1038/ncomms10127.

2. Gebauer, F., Schwarzl, T., Valcárcel, J., and Hentze, M.W. (2021). RNA-binding proteins in human genetic disease. Nat. Rev. Genet. 22, 185–198. 10.1038/s41576-020-00302-y.

3. Greenberg, J.R. (1979). Ultraviolet light-induced crosslinking of mRNA to proteins. Nucleic Acids Res. 6, 715–732.

4. Ule, J., Jensen, K., Mele, A., and Darnell, R.B. (2005). CLIP: A method for identifying protein–RNA interaction sites in living cells. Methods 37, 376–386. 10.1016/j.ymeth.2005.07.018.

5. Ramanathan, M., Porter, D.F., and Khavari, P.A. (2019). Methods to study RNA-protein interactions. Nat. Methods 16, 225–234. 10.1038/s41592-019-0330-1.

6. Patton, R.D., Sanjeev, M., Woodward, L.A., Mabin, J.W., Bundschuh, R., and Singh, G. (2020). Chemical crosslinking enhances RNA immunoprecipitation for efficient identification of binding sites of proteins that photo-crosslink poorly with RNA. RNA 26, 1216–1233. 10.1261/rna.074856.120.

7. Tayri-Wilk, T., Slavin, M., Zamel, J., Blass, A., Cohen, S., Motzik, A., Sun, X., Shalev, D.E., Ram, O., and Kalisman, N. (2020). Mass spectrometry reveals the chemistry of formaldehyde cross-linking in structured proteins. Nat. Commun. 11, 3128. 10.1038/s41467-020-16935-w.

8. Granneman, S., Kudla, G., Petfalski, E., and Tollervey, D. (2009). Identification of protein binding sites on U3 snoRNA and pre-rRNA by UV cross-linking and high-throughput analysis of cDNAs. Proc. Natl. Acad. Sci. U. S. A. 106, 9613–9618. 10.1073/pnas.0901997106.

9. Hafner, M., Katsantoni, M., Köster, T., Marks, J., Mukherjee, J., Staiger, D., Ule, J., and Zavolan, M. (2021). CLIP and complementary methods. Nat. Rev. Methods Primer 1, 1–23. 10.1038/s43586-021-00018-1.

10. Anastasakis, D.G., Jacob, A., Konstantinidou, P., Meguro, K., Claypool, D., Cekan, P., Haase, A.D., and Hafner, M. (2021). A non-radioactive, improved PAR-CLIP and small RNA cDNA library preparation protocol. Nucleic Acids Res. 49, e45. 10.1093/nar/gkab011.

11. Queiroz, R.M.L., Smith, T., Villanueva, E., Marti-Solano, M., Monti, M., Pizzinga, M., Mirea, D.-M., Ramakrishna, M., Harvey, R.F., Dezi, V., et al. (2019). Comprehensive identification of RNA-protein interactions in any organism using orthogonal organic phase separation (OOPS). Nat. Biotechnol. 37, 169–178. 10.1038/s41587-018-0001-2.

12. Shchepachev, V., Bresson, S., Spanos, C., Petfalski, E., Fischer, L., Rappsilber, J., and Tollervey, D. (2019). Defining the RNA interactome by total RNA-associated protein purification. Mol. Syst. Biol. 15, e8689. 10.15252/msb.20188689.

13. Urdaneta, E.C., and Beckmann, B.M. (2020). Fast and unbiased purification of RNA-protein complexes after UV cross-linking. Methods 178, 72–82. 10.1016/j.ymeth.2019.09.013.

14. Esteban-Serna, S., McCaughan, H., and Granneman, S. (2023). Advantages and limitations of UV cross-linking analysis of protein–RNA interactomes in microbes. Mol. Microbiol. 120, 477–489. 10.1111/mmi.15073.

15. Granneman, S., Petfalski, E., and Tollervey, D. (2011). A cluster of ribosome synthesis factors regulate pre-rRNA folding and 5.8S rRNA maturation by the Rat1 exonuclease. EMBO J. 30, 4006–4019. 10.1038/emboj.2011.256.

16. Delan-Forino, C., and Tollervey, D. (2020). Mapping Exosome–Substrate Interactions In Vivo by UV Cross-Linking. In The Eukaryotic RNA Exosome: Methods and Protocols Methods in Molecular Biology., J. LaCava and Š. Vaňáčová, eds. (Springer), pp. 105–126. 10.1007/978-1-4939-9822-7_6.

17. Bresson, S., Shchepachev, V., Spanos, C., Turowski, T.W., Rappsilber, J., and Tollervey, D. (2020). Stress-Induced Translation Inhibition through Rapid Displacement of Scanning Initiation Factors. Mol. Cell 80, 470–484.e8. 10.1016/j.molcel.2020.09.021.

18. van Nues, R., Schweikert, G., de Leau, E., Selega, A., Langford, A., Franklin, R., Iosub, I., Wadsworth, P., Sanguinetti, G., and Granneman, S. (2017). Kinetic CRAC uncovers a role for Nab3 in determining gene expression profiles during stress. Nat. Commun. 8, 12. 10.1038/s41467-017-00025-5.

19. Tufegdzic Vidakovic, A., Harreman, M., Dirac-Svejstrup, A.B., Boeing, S., Roy, A., Encheva, V., Neumann, M., Wilson, M., Snijders, A.P., and Svejstrup, J.Q. (2019). Analysis of RNA polymerase II ubiquitylation and proteasomal degradation. Methods 159–160, 146–156. 10.1016/j.ymeth.2019.02.005.

20. Webb, S., Hector, R.D., Kudla, G., and Granneman, S. (2014). PAR-CLIP data indicate that Nrd1-Nab3-dependent transcription termination regulates expression of hundreds of protein coding genes in yeast. Genome Biol. 15, R8. 10.1186/gb-2014-15-1-r8.

21. Dodt, M., Roehr, J.T., Ahmed, R., and Dieterich, C. (2012). FLEXBAR—Flexible Barcode and Adapter Processing for Next-Generation Sequencing Platforms. Biology 1, 895. 10.3390/biology1030895.

22. Flicek, P., Amode, M.R., Barrell, D., Beal, K., Billis, K., Brent, S., Carvalho-Silva, D., Clapham, P., Coates, G., Fitzgerald, S., et al. (2014). Ensembl 2014. Nucleic Acids Res. 42, D749–D755. 10.1093/nar/gkt1196.

23. Robinson, J.T., Thorvaldsdóttir, H., Winckler, W., Guttman, M., Lander, E.S., Getz, G., and Mesirov, J.P. (2011). Integrative Genomics Viewer. Nat. Biotechnol. 29, 24–26. 10.1038/nbt.1754.

